# Non-specific effect of double-stranded RNAs on Egyptian broomrape (*Phelipanche aegyptiaca*) seed germination

**DOI:** 10.1101/2024.09.19.613516

**Authors:** Nariman Zainali, Houshang Alizadeh, Hassan Alizadeh, Philippe Delavault

## Abstract

Obligate root parasitic plants of the Orobanchaceae family exhibit an intricate germination behavior. The host-dependent germination process of these parasites has prompted extensive research into effective control methods. While the effect of biomaterials such as amino acids and microRNA-encoded peptides have been explored, the effect of double-stranded RNAs (dsRNAs) has remained unexamined during the germination process. In this study, we asked whether an exogenously applied dsRNA can inhibit the germination of a root parasite, P. aegyptiaca. To this end, a dsRNA was designed to target the *CYP707A1* (dsCYP7), a marker gene of the chemically-dependent germination of broomrape seeds. Application of a concentrated dsCYP7 significantly reduced seed germination. However, two non-germination-specific dsRNAs designed to target mannose-6-phosphate reductase and green fluorescent protein brought about similar inhibitions. Moreover, applying rNTPs, which mimic RNA nitrogenous bases, also caused a similar reduction in the germination, suggesting that the non-specific inhibitory effect of the dsRNAs might arise from the nitrogenous moiety thereof. While, dsRNA application inhibited seed germination, their non-specific effects may pose a challenge for their application in studying root parasites germination, emphasizing on the need for further investigations.

## 1 Introduction

The process of seed germination in many orobanchaceous parasitic plants is captivatingly intricate. These parasites produce a huge number of minuscule seeds with long-term viability in the soil (Joel, 2013). In addition, seeds do not germinate unless specific requirements are met. After a so-called conditioning period during which seeds absorb water and get imbibed, they still do not germinate unless they receive a chemical signal showing that an appropriate host plant is in close vicinity (Nelson, 2021). These chemical signals are called germination-stimulant and most of them belong to the strigolactones (SLs) phytohormone family with established roles in shaping plant architecture and its interaction with arbuscular mycorrhizal fungi (Akiyama et al., 2005; Gomez-Roldan et al., 2008; Huizinga and Bouwmeester, 2023). SLs are secreted by the roots of host plants, especially in phosphorus- and nitrogen-depleted conditions (Yoneyama et al., 2007b, 2007a, 2012), and are perceived by parasites via a complex signaling pathway that eventually, breaks the dormancy and leads to seed germination (Brun et al., 2021).

Even though in the past few years our overall knowledge of the molecular events that lead to germination as well as parasitism has been considerably improved, there is still a dire need for an appropriate functional analysis tool (Yoshida and Kee, 2021). In such a situation, RNA interference (RNAi) could help us as an invaluable tool. Being highly specific and the ability to be designed for each gene, it offers a versatile alternative tool not only to study but also to control seed germination in parasitic plants. In the host-parasite interactions, the current view is that the silencing is achievable only after the formation of host-parasite connection via haustorial structure (Jhu et al., 2022) and thus the germination process, or the pre-attachment phases in general, is assumed inaccessible to be studied using this strategy (Yoshida and Kee, 2021). However, there are a few studies showing the functionality of dsRNAs absorbed via seed or root system in silencing their target genes in plants (Jiang et al., 2014; Li et al., 2015; Ludba, 2018). Moreover, a recent new study has opened up a new avenue for environmental RNAi experiments in plants. Betti et al. (2021) showed that the exogenously applied miR399 through the growth medium triggered the silencing of its target gene, *PHO*, in *Arabidopsis thaliana* plants after being absorbed through the root system and movement via the xylem channels. In addition, culturing plants overexpressing the miR399 reduced the expression level of its target gene in the neighboring wild-type *Arabidopsis* plants likely by secreting miRNAs into the environment (Betti et al., 2021). More recently, Tourneur and colleagues (2024) showed that miRNA-encoded peptides (miPEPs) can influence the germination of *Orobanche cumana* when applied exogenously. The researchers used various synthetic miPEPs on the *O. cumana* seeds and found that specific miPEPs significantly inhibited the seed germination by increasing the pri-miRNAs expression and their corresponding target genes downregulation (Tourneur et al., 2024). These lines of evidence may suggest that the process of germination in root holoparasites can be controlled via the manipulation of the genes involved in the germination process by the means of gene silencing (Zainali et al., 2024).

Thus, we asked the question of whether an exogenously applied silencing molecule such as dsRNA can be used to control the seed germination in *P. aegyptiaca*. To answer this question, we selected the *CYP707A1*, an abscisic acid (ABA)-catabolic gene, which has been shown to be up-regulated following treatment with GR24, a synthetic strigolactone analog, and is necessary for the release of dormancy in broomrape species (Lechat et al., 2012; Brun et al., 2019).

## 2 Materials and methods

### 2.1 Plant materials

Seeds of *Phelipanche aegyptiaca* (Pers.) Pomel were collected in September 2020 from infested tomato fields in Sanandaj, Kurdistan (Iran). After sieving through 400, 250, 180, and 165 μm strainers, the fraction between 400-250 μm was collected and stored at 21L until use.

### 2.2 Seed disinfection and conditioning

Seed disinfection was carried out according to Pouvreau et al. (2021). Seeds, in the final density of 10-20 g/L (dry seed weight/v), were added with incubation medium (1 mM HEPES buffer pH 7.5, and 0.1% PPM (v/v)), tubes were wrapped with aluminum foil, and kept in dark at 21 °C for four days before GR24 induction.

### 2.3 Seed and dilution plates preparation

Four days post-conditioning, the incubation media was discarded and equal volumes of dH_2_O and 0.1% sterile agarose were added to set seed density at 10-20 g/L. Afterwards, HEPES and PPM were added as previously described. Finally, seeds were distributed into the 96-well plate (Cell Culture Multiwell Plate Cellstar; Greiner Bio-One) using cut DISTRITIPS® (Gilson, France) in a required volume. Dilution plate was prepared for (+)GR24 containing 10^−5^ to 10^−12^ M concentrations in 1% acetonitrile as described in Pouvreau et al., (2021).

### 2.4 Preparation of dsRNAs

#### 2.4.1 Designing dsRNAs

dsCYP7 was designed as explained in the protocol 1 (Supplementary material, SM). The same procedure was used to design the dsGFP while the same sequence as Farrokhi et al., (2019) was used for dsM6PR.

### 2.4.2 Inducing *PaCYP707A1* expression

Disinfected *P. aegyptiaca* seeds were kept at 21 °C in dark for seven days in the conditioning medium. After seven days, seeds were treated with 10^−6^ M (+)GR24 for maximal germination induction (Yao et al., 2016), and subsequently were incubated at 21 °C in dark up to 18 hours for maximal *CYP707A1* upregulation (Lechat et al., 2012). Afterward, seeds were blotted and dried on tissue paper, transferred into aluminum foil, and immediately snap-frozen in liquid nitrogen.

#### 2.4.3 RNA isolation and cDNA preparation

Seeds were ground to a fine powder in pre-chilled mortars in liquid. RNA was isolated from 100 mg starting seed materials using the NucleoSpin-RNA-Plant kit (MACHEREY-NAGEL, Germany) as per manufacturer’s instructions. The isolated RNAs were treated with RNase-free DNase I set (QIAGENE) for effective removal of DNAs. Samples were then purified using the NucleoSpin RNA Clean-up XS kit (MACHEREY-NAGEL, Germany) and eluted in 20 µL RNase-free H_2_O. The quantity and quality of RNA samples were measured spectrophotometrically and electrophoretically. The first cDNA strand was synthesized from one μg total RNA using the qScript cDNA SuperMix (Quantabio) according to the manufacturer’s instruction.

#### 2.4.4 Amplification of the gene fragments

Two gene specific primers were designed using Primer3 (Untergasser et al., 2012) to amplify the corresponding region of the CYP7 gene fragment (242 bp) and the T7 promoter sequence was added to the 5′ ends of each primer (Table 1). The corresponding fragments for dsM6PR (340 bp) and dsGFP (214 bp) in the L4440 vectors containing them were amplified using the T7 primers presented in Table 1. PCR amplification was carried out in a volume of 100 µL containing 100 µM dNTPs (Promega), 25 µM of each primer, 1 U *Q5* DNA polymerase (New England Biolabs), 2 µL of the first cDNA strand for CYP7, and 10 ng L4440 vectors for M6PR and GFP. Amplification was carried out in a MyCycler thermal cycler (BioRad, USA) with a program included an initial denaturation step at 98 °C for 2 mins, followed by 35 cycles of 10 seconds at 98 °C, 30 seconds at 60 °C, 30 seconds at 72 °C, and ended with a final extension step at 72 °C for 2 mins. Five µL of the amplification reactions was run in the electrophoresis for 30 mins at 100 V in 1% agarose gel. The PCR reaction was purified with the NucleoSpin Gel and PCR Clean-up kit (MACHEREY-NAGEL, Germany) according to the manufacturer’s protocol and the purified fragments were eluted in 20 µL of RNase-free H_2_O. The sequence of the amplified *PaCYP707A1* fragment was also confirmed via Sanger sequencing.

**Table 1.**
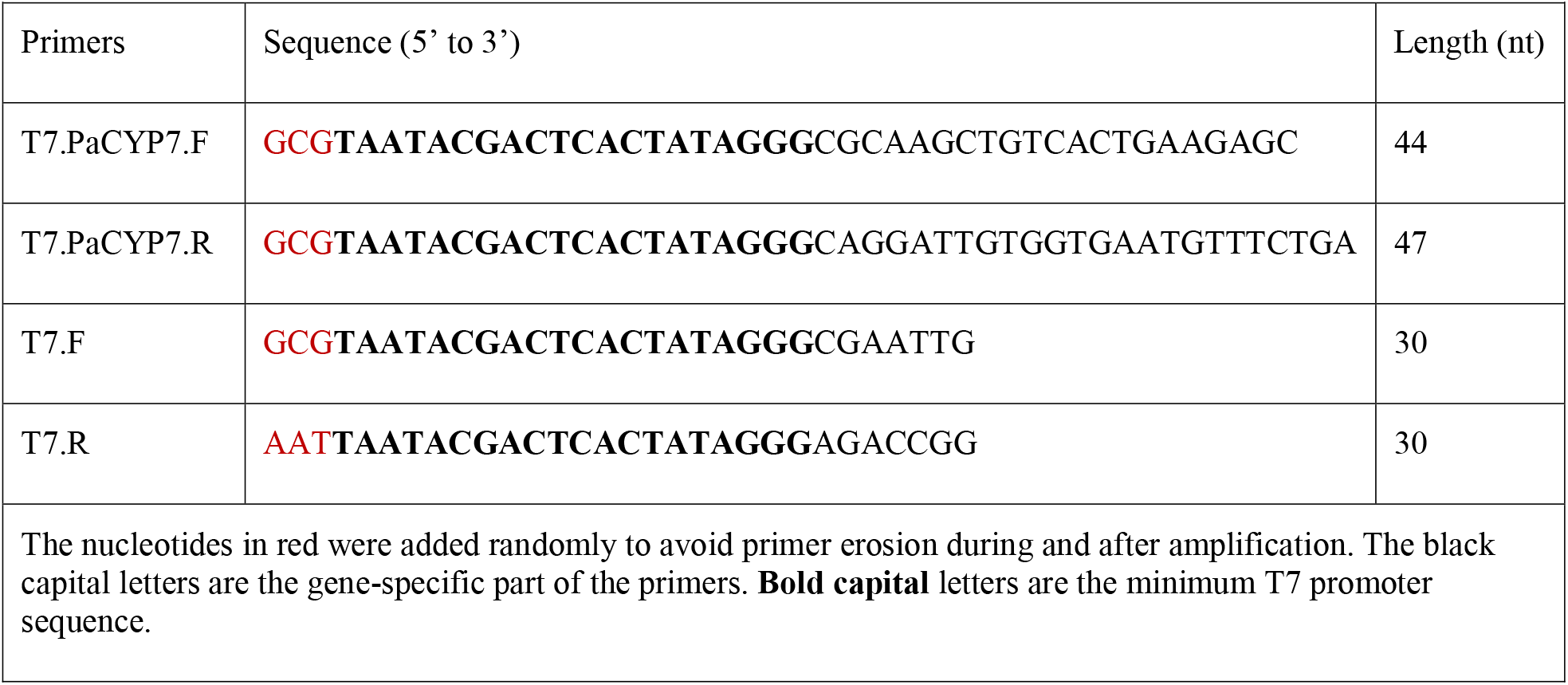
List of the primers used in the current study.

#### 2.4.5 *In vitro* transcription and dsRNA purification

The *in vitro* transcription reactions were performed according to the MEGAscript Kit manual (Invitrogen) in a total volume of 20 µL using 1.5-2 pmoles DNA template by a 16-hours incubation at 37 °C. The *in vitro* transcription reactions were then purified using the MEGAclear kit (Invitrogen) according to the manufacturer’s protocol and the purified dsRNAs were eluted in 100 µL sterile dH_2_O. The quantity and quality of dsRNAs were measured spectrophotometrically and electrophoretically as well. Moreover, the picomolar concentration of the dsRNAs was calculated using the RNA Molecular Weight Calculator tool (https://www.aatbio.com/tools/calculate-RNA-molecular-weight-mw, protocol 2, SM).

#### 2.4.6 Concentrating dsRNAs

To obtain a more concentrated dsRNA, ethanol precipitation with ammonium acetate was performed according to the method presented in MEGAclear kit instruction (Invitrogen). To this end, several purified *in vitro* transcription reactions were pooled and used for precipitation. At the end, the dsRNA pellets were resuspended in a desired volume of sterile dH_2_O.

### 2.5 Treating broomrape seeds with dsRNAs

In the first experiment, dsCYP7 was applied to the final concentration of ∼ 0.7 µM. A dilution plate was prepared as previously described. The seed plate was prepared by adding 40.9 µL seeds (10 g/L) and 4.1 µL dsCYP7 (∼ 8.6 µM) to each well. Afterward, five µL from each (+)GR24 concentration was added to 45 µL seed/dsCYP7 in the corresponding wells of the seed plate producing 10^−6^ to 10^−13^ M (+)GR24 concentration and ∼ 0.7 µM dsCYP7 (Figure S1). In the second experiment, a more concentrated dsCYP7 of ∼ 12 µM was applied on the seeds in the presence of 10^−6^-10^−8^ M (+)GR24 (Figure S2).

However, in the subsequent experiments, three concentrations of ∼ 6, 3, and 1.5 µM was applied for dsCYP7, dsM6PR, and dsGFP (Figure S3-4). In addition, rNTPs (MEGAscript kit, Invitrogen) was used in three concentrations of ∼ 5, 2.5, and 1.25 mM (protocol 3, SM, Figure S4). It is worth noting that in these experiments, 10^−6^ M (+)GR24 was applied for germination induction and no serial dilution of (+)GR24 was used. After treatment, the seed plates were kept at 21 °C in dark for four days.

### 2.6 Staining and Absorbance Reading

In the first experiment with 0.7 µM dsCYP7, seeds were added with 5 µL Thiazolyl Blue Tetrazolium Bromide (MTT, 5 g/L, SIGMA-Aldrich) per well eight days-post treatment and kept at 21 °C in dark for one day. The day after, 100 µL of solubilization buffer (10% Triton X-100 and 0.04 M HCl in isopropanol) was added per well.

### 2.7 Data collection and analysis

The seed plates were regularly monitored from day one to eight after treatment under a binocular (Olympus SZX10; Olympus Europa GmbH) and the number of germinated seeds was determined. The seeds with protruded radicals were considered as germinated throughout our experiments. The relative germination ratio of each well was calculated as below:

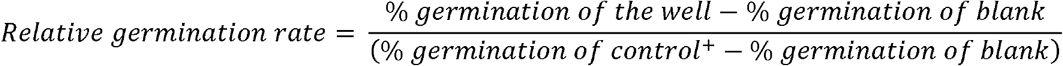

where control+ was the germination of the wells containing 10^−6^ M (+)GR24 and blank was the germination of the wells without (+)GR24.

In the experiment with 0.7 µM dsCYP7, 570 and 630 absorbances were read with a Polarstar Omega using an UV/Vis spectrometer (BMG Labtech) after MTT staining and the dose-response curve was obtained as described previously (Pouvreau et al., 2021) using the *drc* package V3.0.1 (Ritz et al., 2015) in R V4.3 (R Core Team, 2023). In this experiment, the absorbances of the wells were used to obtain the dose-response curve. However, in the experiment with 12 µM dsCYP7, the relative germinations rates were directly calculated from the germination rate of the wells and were used to obtain the dose-response curve because the wells containing dsCYP7 showed background color making absorbance reading impossible. In the rest of the experiments, the relative germination was also obtained from the germination rate of the wells as well.

The Kruskal-Wallis non-parametric test was used to infer the statistical differences among the treatments as the calculated relative germination ratios were not normally distributed, and the multiple pairwise comparisons were performed using the Dunn’s Test with Bonferroni correction for *p-values* in Minitab V21.4.2. Data were visualized using *ggplot2* package V3.5.0 (Wickham, 2016) in R V4.3 (R Core Team, 2023).

## 3 Results

### 3.1 Amplification of the *PaCYP707A1* gene fragment and dsCYP7 preparation

The result of BLASTp for the *PaCYP707A1* sequence retrieved from the PPGP database showed the ABA 8’-hydroxylase protein in *P. ramosa* as the best hit indicating that the sequence corresponded to the same gene in *P. aegyptiaca*. Moreover, Sanger sequencing showed that the amplified *PaCYP707A1* fragment was almost identical to the sequence retrieved from the PPGP database (Figure S5**Error! Reference source not found**.). The agarose gel electrophoresis confirmed the production of the corresponding products with their expected sizes of ∼290 and ∼240 bp for *PaCYP707A1* PCR product and dsCYP7, respectively. Considering that *PaCYP707A1* fragment contained two T7 promoter sequences, its size was larger than that of dsCYP7 (Figure S6).

### 3.2 Exogenous application of dsRNAs on *P. aegyptiaca* seeds

#### 3.2.1 dsCYP7 treatment in a gradient of (+)GR24

Initially, dsCYP7 was applied to the seeds at the final concentration of 0.7 µM in a gradient of (+)GR24 (Figure S1). The seed plate was monitored regularly from the day one to eight after treatment. Staining with MTT and germination bioassays were conducted to check the seed viability state eight days post treatment (8-dpt). No significant changes in the relative absorbances were observed in the seed treated with 0.7 µM dsCYP7 compared with seeds treated only with (+)GR24 up to 8-dpt (*p-value* > 0.05, Figure 1A, Figure S7) indicating that dsCYP7 had no or negligible impact on the seed germination at the applied concentration.

**Figure 1.**
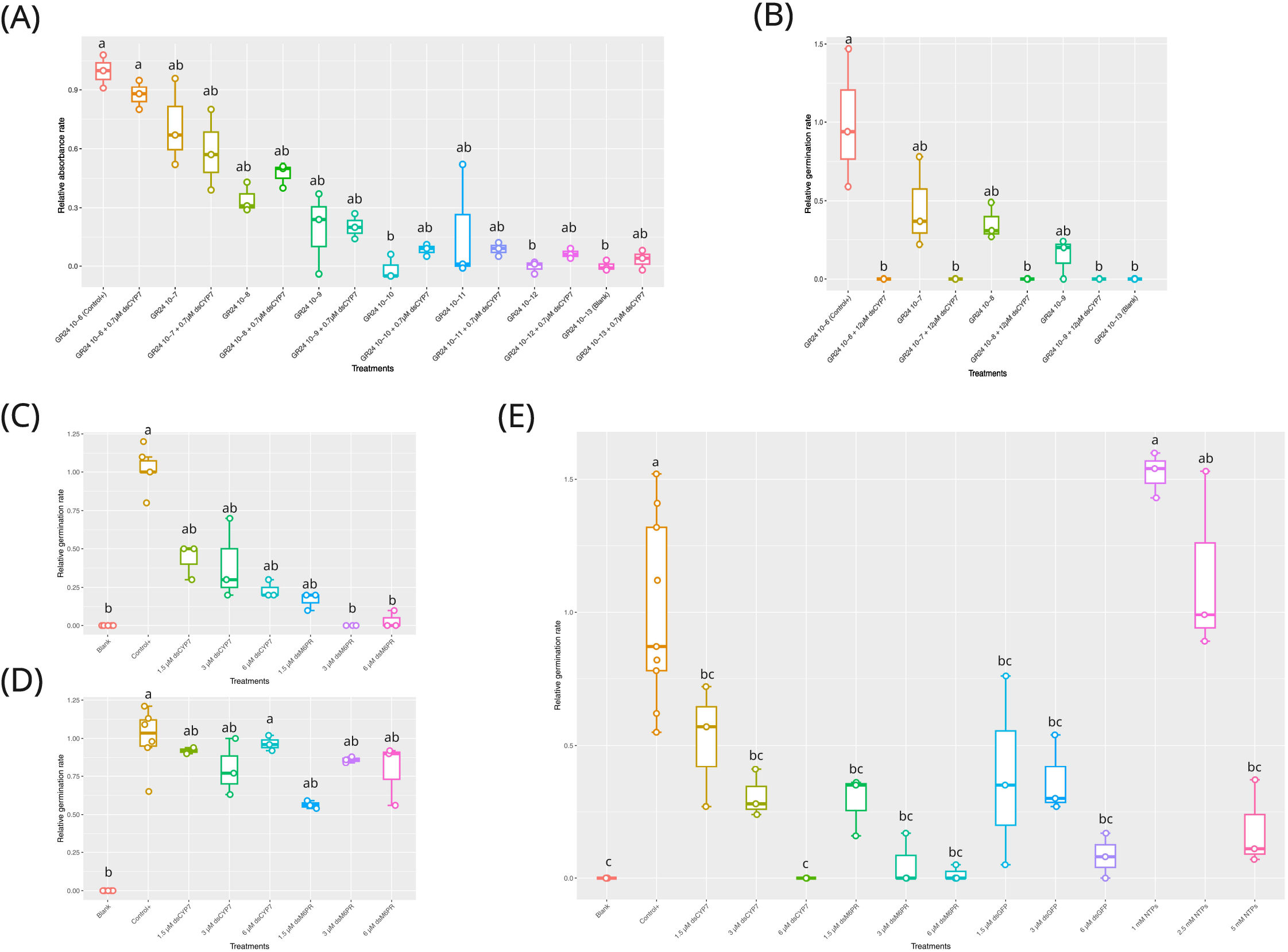
The effect of different dsRNA treatments on the germination of Egyptian broomrape, *P. aegyptiaca*. **(A)** The boxplot for the relative absorbance rates for the experiment with 0.7 µM dsCYP7. 1o^-6^ M +GR24 was considered as the positive control and seeds treated with 10^−13^ M +GR24 were considered as blank. **(B)** The boxplot for the relative germination rates for the experiment with 12 µM dsCYP7. (C) The boxplot of the relative germination rates obtained in the experiment with dsCYP7 as well as dsM6PR as a non-specific dsRNA on the eitghth day (8-dpt). Seeds were treated with 6, 3, and 1.5 µM of each dsRNA. Seeds without +GR24 were considered as blank. **(D)** The relative germination rates of the seeds in the experiment presented in C, eight days after washing. **(E)** The box plot of the relative germination rates in the experiment with dsGFP and rNTPs at 7-dpt. dsGFP was applied at 6, 3, and 1.5 µM concentration while rNTPs was applied at 5, 2.5, and 1.25 mM concentrations. The statistical differences among treatments were determined using the Kruskal-Wallis non-parametric test and multiple comparisons with the Dunn’s test and Bonferroni correction for p-values. Treatments sharing a same lowercase letters are not significantly different *(p-value* > 0.05).

Therefore, a more concentrated dsCYP7 at the final concentration of ∼ 12 µM was applied on the seeds in the presence of 10^−6^-10^−8^ M (+)GR24 (Figure S2). Interestingly, no germinated seed was observed in the wells with 12 µM dsCYP7 up to 7-dpt. Because the dsCYP7-treated seed wells showed background color after staining with MTT, most likely due to a reaction between dsRNAs and the components of the solubilization solution, relative absorbances were not informative and absorbance reading was not able to produce a correct dose-response curve reflecting the effect of the dsCYP7 (data not shown). The germination rate of the dsCYP7-treated seeds at 8-dpt reduced significantly compared to the positive control (*p-value* < 0.001) suggesting an inhibitory effect for dsCYP7 on the broomrape germination (Figure 1B).

#### 3.2.2 dsCYP7 alongside negative controls

In order to examine the reproducibility of the previous experiment and to test the specificity of the dsCYP7 in inhibiting broomrape germination, another experiment was carried out by including a dsRNA targeting M6PR gene (dsM6PR, Figure S8 A & B) as a gene without an apparent role in the seed germination. In this experiment, three different concentrations of each dsRNAs e.g. 6, 3, and 1.5 µM were applied in the 10^−6^ M concentration of (+)GR24. Unexpectedly, a reduction in the seed germination was recorded in the seeds treated with both dsRNAs compared to the untreated seeds (Figure 1C, Figure S9). In addition, the germination rates of the seeds treated with 6 and 3 µM dsM6PR was significantly lower compared to those of the positive control at 7-dpt (*p-value* < 0.001) (Figure 1C). Although the effects of dsRNAs on germination rates demonstrated a clear dose-dependent relationship, the differences among dsRNAs concentrations were not statistically significant (*p-value* > 0.05) (Figure 1C). On average, germination rates were reduced approximately 79%, 62%, and 65% in the seeds treated with 6, 3, and 1.5 µM of dsCYP7, respectively, compared to the positive control. In addition, seeds treated with the same concentrations of dsM6PR exhibited reductions of 100%, 100%, and 85%, respectively.

To rule out that the lack of germination in the dsRNA-treated seeds was not due to the toxicity of the dsRNAs, a washing step was carried out at 8-dpt. Eight days after washing, the germination rate of the dsRNA-treated seeds recovered to the level of the positive control (*p-value* > 0.05) (Figure 1D) indicating that dsRNAs were not toxic. It is also worth noting that the removed dsCYP7 and dsM6PR from the wells of the seed plate during the washing step, were precipitated and purified to check whether the applied dsRNAs were degraded following seed treatment or remained intact until the end of the experiments. Agarose gel electrophoresis showed that the most of the applied dsRNAs remained intact (Figure S10).

Unexpectedly, dsM6PR caused a more reduction in the seed germination compared to the dsCYP7. It was not clear that this reduction in the germination arose from an unknown role of the M6PR gene and caused by M6PR gene silencing or from an unknown non-specific effect of the dsRNAs on the germination. To understand it better, another experiment was conducted by including a dsRNA designed to target the green fluorescent protein (GFP) sequence (dsGFP, Figure S8 C & D). Moreover, ribonucleotides (rNTPs) at 5, 2.5, and 1.25 mM concentrations were included in the experiment as another control. Similar to the previous experiments, the germination of broomrape was considerably reduced in the wells treated with dsCYP7 and dsM6PR, almost at a similar level (Figure 1E). Moreover, a similar pattern of germination inhibition was also recorded in the seeds treated with different concentrations of dsGFP and the germination was significantly reduced compared to the positive control (*p-value* < 0.001) (Figure 1E). While no germination inhibition was observed in the seeds treated with 1.25 and 2.5 mM rNTPs compared to the positive control, the germination was significantly reduced in the seeds treated with 5 mM rNTPs compared to the positive control (*p-value* < 0.001), but it was not significantly different from the dsRNA-treated seeds (*p-value* > 0.05) (Figure 1E). In fact, the germination of the seeds treated with 1.25 and 2.5 mM rNTPs were increased, on average, 52% and 14% respectively, compared to the positive control even though the differences among them were insignificant (*p-value* > 0.05). On the other hand, on average, the germination rates were reduced 100%, 92%, 98%, and 63% in the seeds treated with 6 µM dsCYP7, dsM6PR, and dsGFP, as well as 5 mM rNTPs, respectively.

## 4 Discussion

Root parasitic plants belonging to the Orobanchaceae family exhibit a captivatingly intricate germination behavior attracting many researchers all over the world. Even though it is one of the most studied aspects of holoparasites lifecycle, our understandings of the process at the molecular level is still incomplete. Moreover, considering the dependency of the root holoparasites on their host plants that are mostly crop species, the germination is a vital step, making it the focus of several studies in order to find a solution that has an edge over the chemicals to control broomrapes by preventing germination or by enhancing it!

In the current experiment, we asked the question of whether a dsRNA can be used to modulate *P. aegyptiaca* seed germination by targeting the genes involved in the germination process. To this end, *PaCYP707A1* gene was selected as an appropriate target as it has been shown to be involved in the dormancy breaking of broomrape species (Lechat et al., 2012; Brun et al., 2019). The Initial concentration of 0.7 µM dsCYP7 was applied on the seeds and the monitoring was continued up to, usually, 7-dpt as no changes in germination is expected after this time (Pouvreau et al., 2013). Due the lack of noticeable effect at this concentration, a more concentrated dsCYP7 of ∼ 12 µM was applied on seeds as an extremum amount, which completely inhibited the seed germination implying on the effectiveness of dsCYP7. Unexpectedly, however, the application of two non-specific control dsRNAs targeting genes without roles in the germination process, namely dsM6PR and dsGFP, brought about significant reductions in the seed germination in a dose-dependent manner, a similar pattern observed for dsCYP7 indicating that the detected inhibitory effect of the dsCYP7 was likely not a specific effect on the *PaCYP707A1* mRNA level. Moreover, recovery of the germination in the washed dsRNA-treated seeds to a level similar to the positive control indicated that the dsRNAs were not toxic for the seeds and seeds were not killed by dsRNA treatments.

Since both specific and non-specific dsRNAs led to significant reductions in the seed germination of broomrape, we also asked whether the lack of germination in the dsRNA-treated seeds could arise from the effect of the nitrogenous nature of dsRNAs. To address this question, rNTPs were included in the experiment as a control to test the effect of nitrogen bases belonging to RNAs. An inhibition of seed germination at 5 mM concentration of rNTP was achieved, a similar molar concentration to the 6 µM dsRNAs considering their sizes. While the germination rate of the seeds treated with 5 mM rNTPs were, on average, more than 30% higher than that of obtained from the seeds treated with 6 µM dsRNAs, they were not statistically different from each other. This may hint that the inhibitory effect of dsRNAs might arise from the nitrogenous nature thereof. However, the germination rates of the seeds treated with the lower concentrations of rNTPs not only were not reduced, but also, apparently, were enhanced compared to the positive control, an aspect that in turn worths deeper attention and further investigations.

It has been reported that nitrogen deficiency can induce the production and secretion of a strigolactone in sorghum, a host for the root hemiparasitic plant *Striga hermonthica* (Yoneyama et al., 2007a), as well as other plant species (Yoneyama et al., 2012). On the other hand, nitrogenous compounds and nitrogen fertilizers have been shown to reduce the germination and radical elongation in *Striga* and broomrape species (Farnia et al., 1985; Pieterse, 1991; Westwood and Foy, 1999), and replacing the conditioning solution containing nitrogen with a solution without it can neutralize the inhibitory effect of the nitrogen after a few days (Van Hezewijk and Verkleij, 1996), a similar result that observed in the current experiment. Why and how nitrogen inhibits the seed germination in parasitic plants, at molecular and mechanistic levels, still remains to be seen. Recently, it has been shown that ammonium ions can impair the root chemotropism toward SLs in *Phtheirospermum japonicum* likely by regulating the expression of an auxin efflux transporter, *PIN2*, in the roots (Ogawa et al., 2022). Moreover, nitrogen has been shown to repress the formation and growth of haustoria in *P. japonicum* by increasing the content of ABA (Kokla et al., 2022). These lines of reports may suggest that root parasitic plants have evolved a mechanism to perceive information on the state of their rhizosphere, such as the amount of nitrogen, to leverage their germination in the finest situations fitting their heterotrophic nature.

Several studies have investigated the effect of amino acids on the broomrape germination and some have been shown to reduce broomrapes germination when applied exogenously on the seeds (Vurro et al., 2006; Fernández-Aparicio et al., 2017; Arslan et al., 2023; Gibot-Leclerc et al., 2024). While, the effects of amino acids have been suggested to be related to the interference with the biosynthesis pathways of other amino acids (Vurro et al., 2006), whether the observed inhibitory effect reported for them in the aforementioned studies has any link with the nitrogenous moiety thereof, worth further investigations. Furthermore, tryptone, as the main source providing nitrogen in broth media, has been reported to reduce broomrape germination in a dose-dependent manner most likely due to its _L_-tryptophan (Trp) fraction (Kuruma et al., 2021). More recently, it has been shown that exogenously applied Trp is converted into indole-3-acetic acid (IAA) in *O. minor* seeds, which in turn prevents radical elongation (Tsuzuki et al., 2024). It is also worth investigating the possible links between nitrogen and the inhibitory effect of IAA in the seeds of root parasites as the exogenous nitrogen may provide a source for the production of amino acids such as Trp.

Modulating the broomrape seed germination has been achieved via the application of miPEPs by regulating their corresponding *MIR* genes, and subsequently, post-transcriptional gene silencing of their corresponding target genes (Tourneur et al., 2024). miPEPs act specifically by increasing their corresponding *MIR* genes transcription (Lauressergues et al., 2015), leading to germination inhibition when applied exogenously on the broomrape seeds at appropriate amounts (Tourneur et al., 2024). Therefore, the process of germination seems accessible to be modulated through RNA silencing via the exogenous application of molecules such as si/miRNAs and dsRNAs. However, the application of dsRNAs will require refinements as they impacted the phenotype of germination in a non-specific manner in our study. As it seemed that the observed inhibitory effects of dsRNAs on the germination was the result of their nitrogenous nature, the application of exogenous RNAi for studying germination may confront a challenge as the triggering molecules of RNAi all are the same in nature.

## 5 Authors contributions

PD and NZ conceived the idea. NZ carried out the experiments, collected data and analyzed them. PD and HoA provided funding for the experiment. NZ wrote the first draft of the manuscript. PD, HoA, and HaA critically reviewed the manuscript. All authors read and agreed on the final version of the paper.

## 7 Acknowledgement

Mojgan Gholami Malekroudi is thanked for her assistances during the course of the experiments.

## 9 Data Availability

Data of the current study would be available upon reasonable request to the corresponding authors.

