## Supplementary Materials for "Non-specific effect of double-stranded RNAs on Egyptian broomrape (*Phelipanche aegyptiaca*) seed germination"

Supplementary Material

### 1 Protocol 1. Designing dsCYP7

The mRNA sequence of *CYP707A1* from *P. ramosa* (*PrCYP707A1*, JQ838174.1) was BLASTed against *P. aegyptiaca*’s transcriptome, PhAeBC6.final, from PPGP (Yang et al., 2015), using the BLAST feature (Altschul et al., 1990) included in the BioEdit software v7.2. (Hall, 1999). The whole sequence of the first hit (ID: PhAeBC6_h_c21860_g10399_i2) was then searched for ORFs using NCBI orffinder (<https://www.ncbi.nlm.nih.gov/orffinder/>). The longest found ORF was then BLASTed against the “UniProtKB/Swiss-Prot” database and with “higher plants” as organism using BLASTp. The PhAeBC6_h_c21860_g10399_i2 contig was aligned to *PrCYP707A1* complete mRNA seq with ClustalW (Thompson et al., 1994) to determine the complete mRNA sequence of *PaCYP707A1.* The whole aligned sequence of PhAeBC6_h_c21860_g10399_i2 to *PrCYP707A1* (1768 nt) was then considered as the complete *PaCYP707A1* mRNA and used for dsRNA design. Designing the dsCYP7 was done using *PaCYP707A1* mRNA seq as a query by the *pssRNAit* webserver (Ahmed et al., 2020) with the default parameters. Afterwards, the parameters for siRNAs and dsRNA design were adjusted to have minimum off-targets against *S. lycopersicum* as presented in Table S1. The suggested dsRNA by the *pssRNAit* webserver was used in the current study (Table S2).

| Table S1. Parameters for pssRNAit | | | |
| --- | --- | --- | --- |
| Parameters for siRNA design: | | | |
| siRNA Efficiency: 9.0  Range: 0-10, the more the better | Target accessibility (UPE): 15  Range: 0-25, the less the better | | Max # of off-target: 0 |
| Parameters for off-target analysis using psRNATarget: | | | |
| Expect: 0.0  Range 0-5, the less the less off-targets | | Off-target Accessibility (UPE): 0.0  Range: 0-25, the less the less off-targets | |
| Parameters for VIGS candidates design: | | | |
| Range of VIGS length: 100 to 300 | Minimal # of siRNAs in VIGS candidates: 10 | | Minimal distance of two effective siRNAs: 10 |

| Table S2. The suggested dsRNA for *PaCYP707A1* gene by the pssRNAit webserver and the features of the designed dsRNA. | | | | | |
| --- | --- | --- | --- | --- | --- |
| Range on target sequence | Length | # of siRNAs | siRNA sequences | # of off-target | Significant off-targets / # of hits |
| 959-1133 | 175 | 10 | UUGCUCUUCAGUGACAGCUUG  UAUCAAAGCUUCUUGCUCUUC  UUAGACUUGUGUUUUCACCUU  AUCUUCUUUGUGUCUGCCCAA  UUGUCAUAGGCAUCUUCUUUG  UUGAAUAACCCUUGUUGUCAU  AUAGAGGCAGCUCUAAGUGUU  UAAAAGACAAAAUAGAGGCAG  UCAACAGCUUCUCUGAAAGUA  UAAACUCAACAUCUUCAACAG | 0 | - |

The same procedure was used to design dsGFP while the sequence for dsM6PR was adapted from (Farrokhi et al., 2019).

### 2 Protocol 2. Calculating the molar concentration of dsRNAs to apply on the broomrape seeds.

dsCYP7 was purified using the MEGAclear kit (Invitrogen). The concentration of one purified dsCYP7 was **1335.9** ng/µL.

|  | Length* (bp) | Concentration (ng/µl) | The total amount of dsRNA (µg) |
| --- | --- | --- | --- |
| dsCYP7 | 242 | 1336 | 133.6 |
| * The length of transcript after transcription by T7 RNA polymerase. | | | |

The molar concentration of dsCYP7 was calculated as follows as described in Wise et al. (2022). The molecular weight (MW) of *PaCYP707A1* ssRNA was estimated using the RNA Molecular Weight Calculator (https://www.aatbio.com/tools/calculate-RNA-molecular-weight-mw):

- PaCYP7A1 sense RNA seq:
- GCAAGCUGUCACUGAAGAGCAAGAAGCUUUGAUAAAAUCAAAAGGUGAAAACACAAGUCUAAAUUGGGCAGACACAAAGAAGAUGCCUAUGACAACAAGGGUUAUUCAAGAAACACUUAGAGCUGCCUCUAUUUUGUCUUUUACUUUCAGAGAAGCUGUUGAAGAUGUUGAGUUUAAUGGAUAUCUCAUUCCAAAAGGAUGGAAAGUUUUACCACUUUUCAGAAACAUUCACCACAAUCCUG
- MW of *PaCYP7* sense RNA strand: 77.91 kDa
- MW of *PaCYP7* antisense RNA strand: 77.24 kDa
- MW of dsCYP7 = 77.91 kDa + 77.24 kDa = 155.15 kDa or kg/mol
- $dsCYP7 molar concentration=\frac{1335.9\times{10}^{-9}g}{155.15\times{10}^{3}g}=8.61\times{10}^{-12}mol=8.61\frac{pmol}{\mu L}=8.61 \mu M$

The same procedure was used to estimate the concentration of dsM6PR and dsGFP based on their sequences (Below):

- PaM6PR sense RNA seq (390 bp, the **bold** nucleotides belong to the L4440 vector backbone):
- **GCGAAUUGGGUACC**UUGAGGUUGGAAGAGGAACAAUACAGUUUUAGUUAAGUUCUUAAGUAGAUGAAAUAUUGCGAGUUUUUGCCCCACUGCAAUAAUAAACACUAUUUUACCAUAGAAUGGCACACUCCUUUUCCGUCUCUCUCUCUCUCUCUCCUUGUGUUGCAUCAUACAUGAAUGAAAAUCCCUAACAUGGGAGAGAAACUUAUGCGAAAAGAUCGAUACCCCAGAACUUGGCAGGUUGAUUAGUUCUGUAUUUCCGCUCCAUAGUCUUCAACAGUUCCAUAUCCUCAUUGGAGAGCUCGAAGUCAAAAACUUGAAAAUUCUCGAUCAAUCUGUCCUGUUUUGAUGUCUUGGG**GAAUUCAUCGAUGAUAUCAGAUCUGCCGGUCUC**
- MW of sense *PaM6PR* RNA strand: 124.68 kDa
- MW of antisense *PaM6PR* RNA strand: 125.36 kDa
- MW of dsM6PR = 124.68 kDa +125.36 kDa = 250.04 kDa or kg/mol
- GFP sense RNA seq (260 bp, the bold nucleotides belong to the L4440 vector backbone):

**GCGAAUUGGGUACC**UCCUUGAAGUCGAUGCCCUUCAGCUCGAUGCGGUUCACCAGGGUGUCGCCCUCGAACUUCACCUCGGCGCGGGUCUUGUAGUUGCCGUCGUCCUUGAAGAAGAUGGUGCGCUCCUGGACGUAGCCUUCGGGCAUGGCGGACUUGAAGAAGUCGUGCUGCUUCAUGUGGUCGGGGUAGCGGCUGAAGCACUGCACGCCGUAGGUGAAGGUGGUCA**GAAUUCAUCGAUGAUAUCAGAUCUGCCGGUCUC**

- MW of GFP sense RNA strand: 84.1 kDa
- MW of GFP antisense RNA strand: 83.88 kDa
- MW of dsGFP = 84.1 kDa + 83.88 kDa = 168.98 kDa or kg/mol

If required, several *in vitro* transcribed dsRNAs were pooled, ethanol-precipitated, and resuspended in a smaller volume of sterile dH_2_O to obtain a more concentrated dsRNA solution. Finally, dsRNAs were applied on the seeds as shown in Figures S1-4.

### 3 Protocol 3. Calculating rNTPs concentrations

In order to understand whether the inhibition of the seed’s germination was because of the nonspecific influence of the rNTPs of the dsRNA strands rather than the specific effect of the dsRNAs, rNTPs were used as a control. To this end, rNTPs was needed to be applied in an appropriate amount. Thus, we tried to have, almost, the same amount of rNTPs that have been applied through dsRNAs. All dsRNAs, including dsCYP7, dsM6PR and dsGFP, were applied at the concentrations of ~ 6, 3, and 1.5 µM. Therefore, the concentration of rNTPs in the case of the applied dsRNAs can be calculated by multiply the length of the each dsRNA by its concentration. In addition, the lengths are expressed as base pairs (bp) and therefore, it must be considered as well. Consequently:

- Molar concentration of rNTPs = The molar concentration of dsRNA × length × 2
- dsCYP7 = 6.5 × 242 × 2 = 3146 µM = 3.146 mM
- dsM6PR = 6.5 × 390 × 2 = 5070 µM = 5.07 mM
- dsGFP = 6.5 × 261 × 2 = 3393 µM = 3.393 mM

The highest rNTPs concentration was calculated to be about 5 mM in the case of dsM6PR. Thus, this concentration and 2, and 4-fold dilutions thereof were selected to be used as controls.

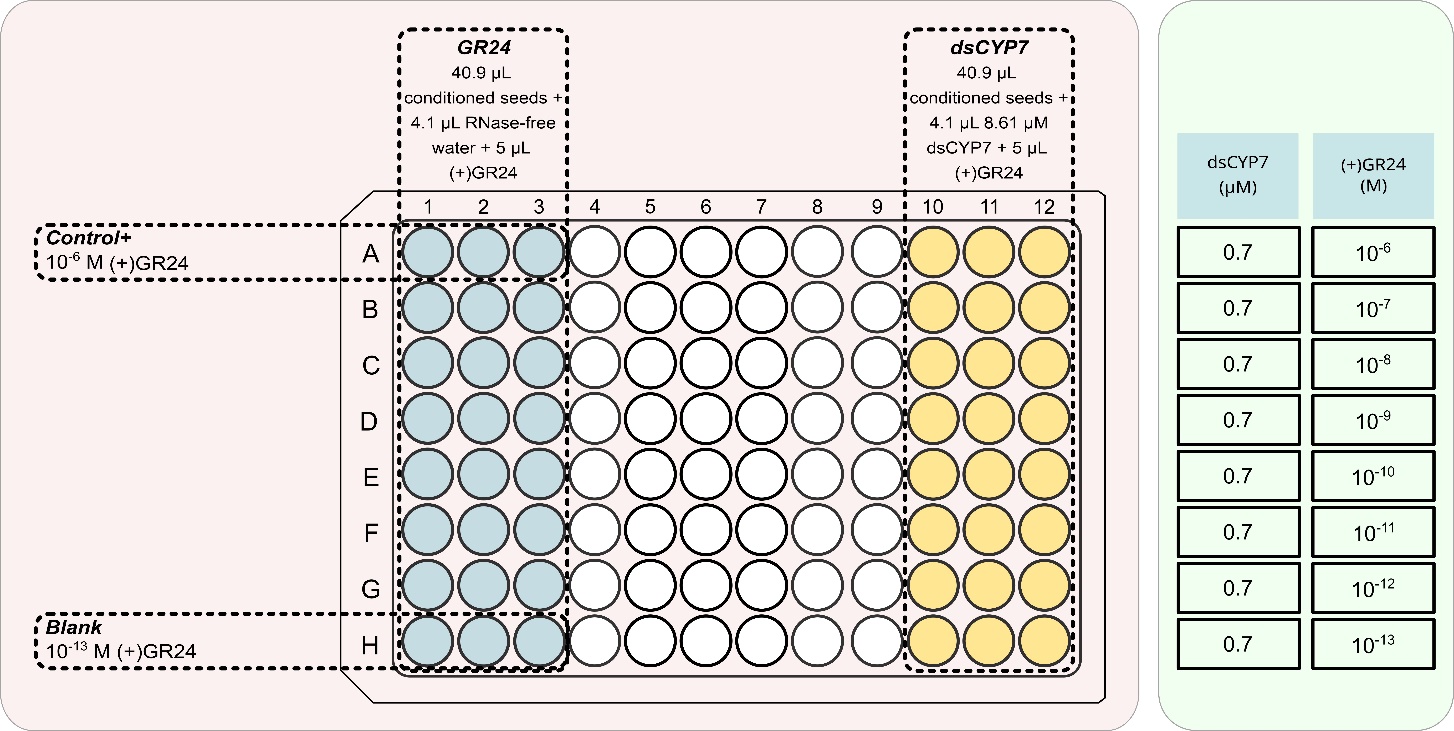

Figure S1. The layout of the seed plate for applying dsCYP7 on broomrape seeds. dsCYP7 was added in the corresponding wells at the final concentration of 0.7 µM in a gradient of (+)GR24. 10^-6^ M (+)GR24 was considered as the positive control while 10^-13^ M (+)GR24 was considered as the negative control (blank).

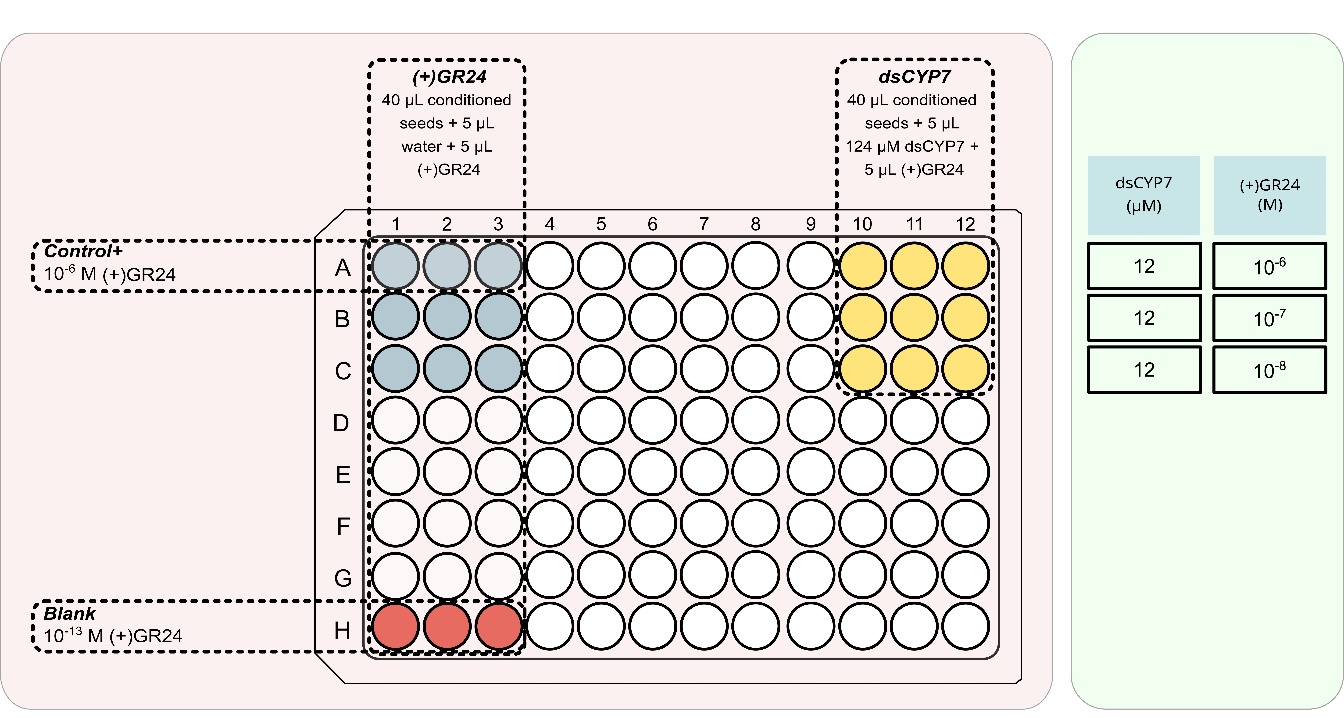

Figure S2. The seed plate layout in the experiment with 12 µM dsCYP7. dsCYP7 was added in the corresponding wells at the final concentration of 12 µM. 10^-6^ M (+)GR24 was considered as the positive control while 10^-13^ M (+)GR24 was considered as the negative control (blank).

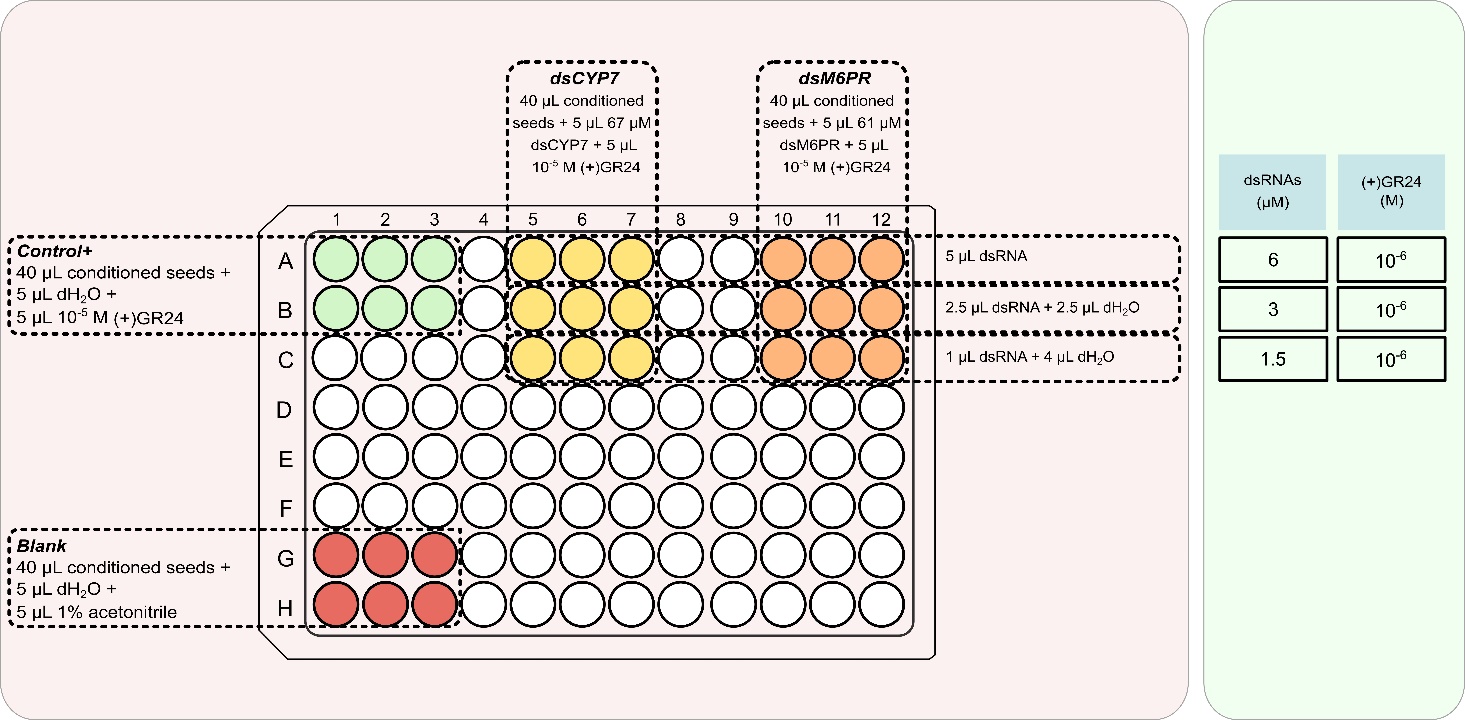

Figure S3. The seed plate layout in the experiment with dsCYP7 and dsM6PR. Both dsRNAs were added in the corresponding wells at three concentrations of 6, 3, and 1.5 µM. All dsRNA-treated wells contained 10^-6^ M (+)GR24 for germination induction. 10^-6^ M (+)GR24 was considered as the positive control while wells without (+)GR24 treatment were considered as the negative control (blank).

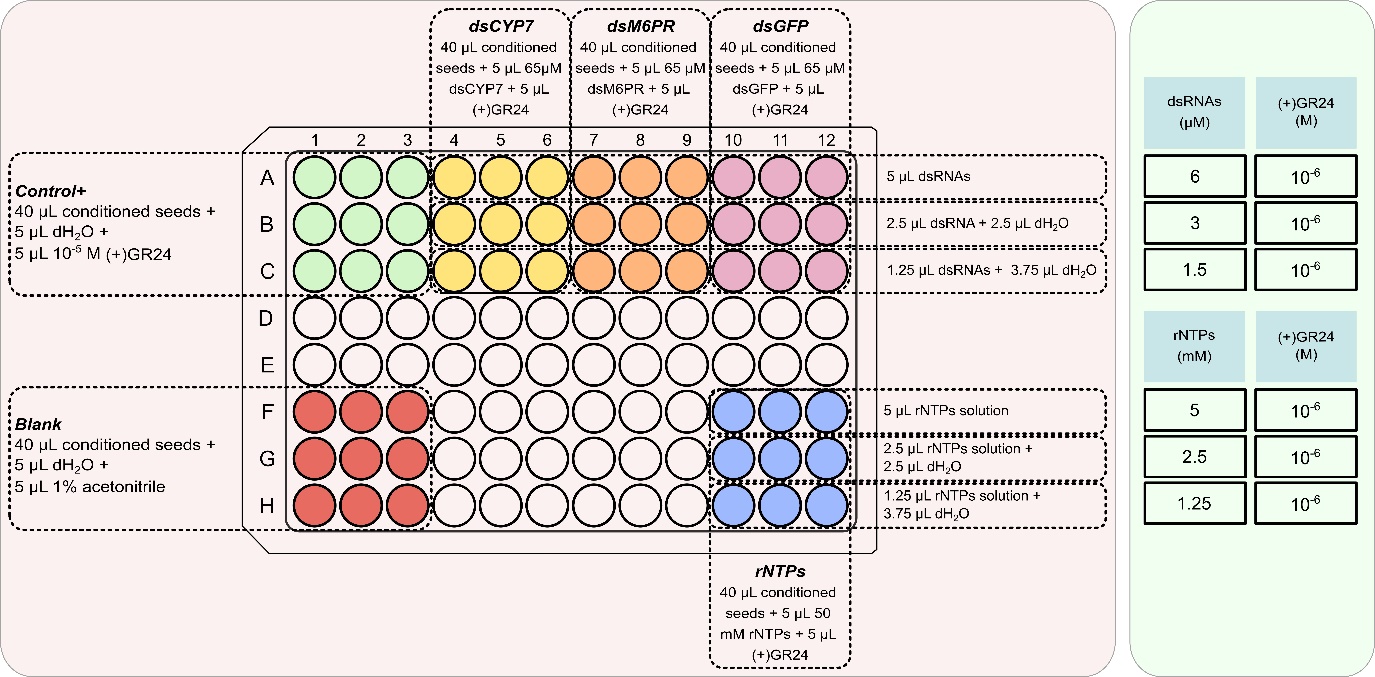

Figure S4. The seed plate layout in the experiment with dsGFP and rNTPs. All dsRNAs were added in the corresponding wells at three concentrations of 6, 3, and 1.5 µM. rNTPs was used at three concentration of 5, 2.5, and 1.25 mM. All dsRNA- and rNTP-treated wells contained 10^-6^ M (+)GR24 for germination induction. 10^-6^ M (+)GR24 was considered as the positive control while wells without (+)GR24 treatment were considered as the negative control (blank).

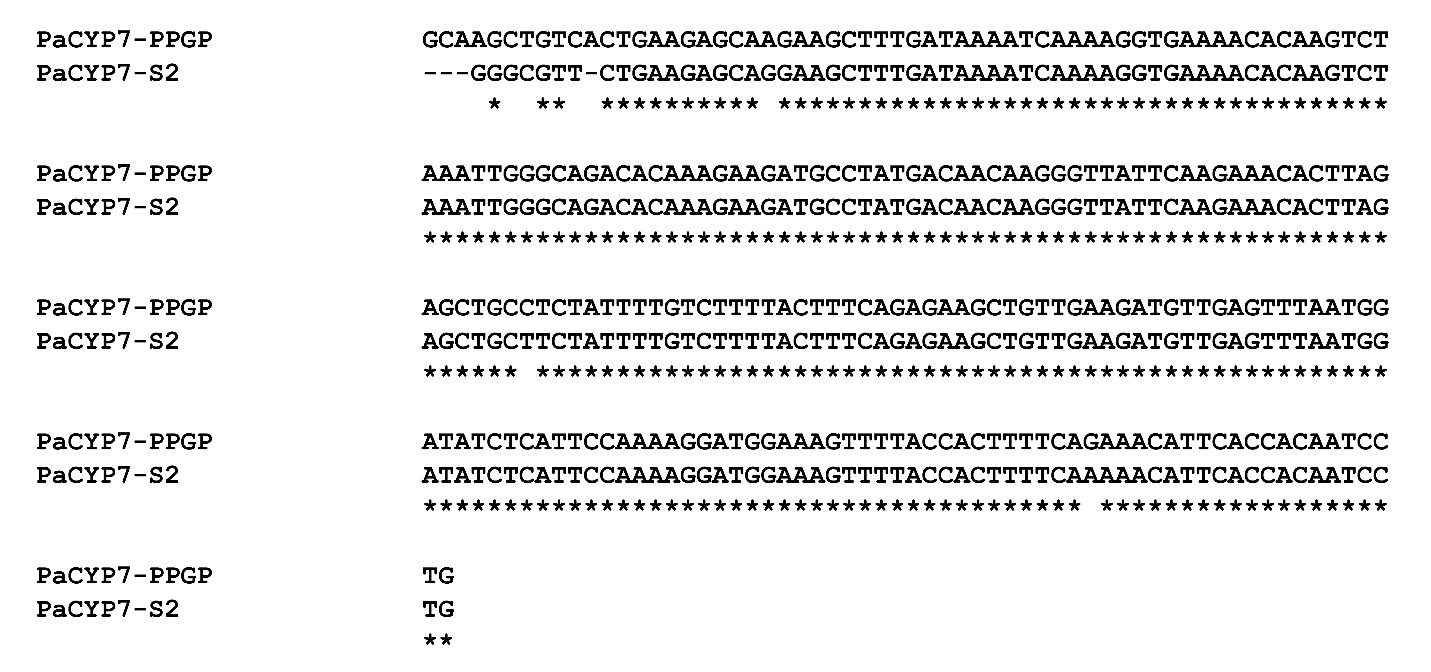

Figure S5. Alignment of the amplified fragment of *PaCYP7A1* against the same sequence obtained from the PPGP transcriptome.

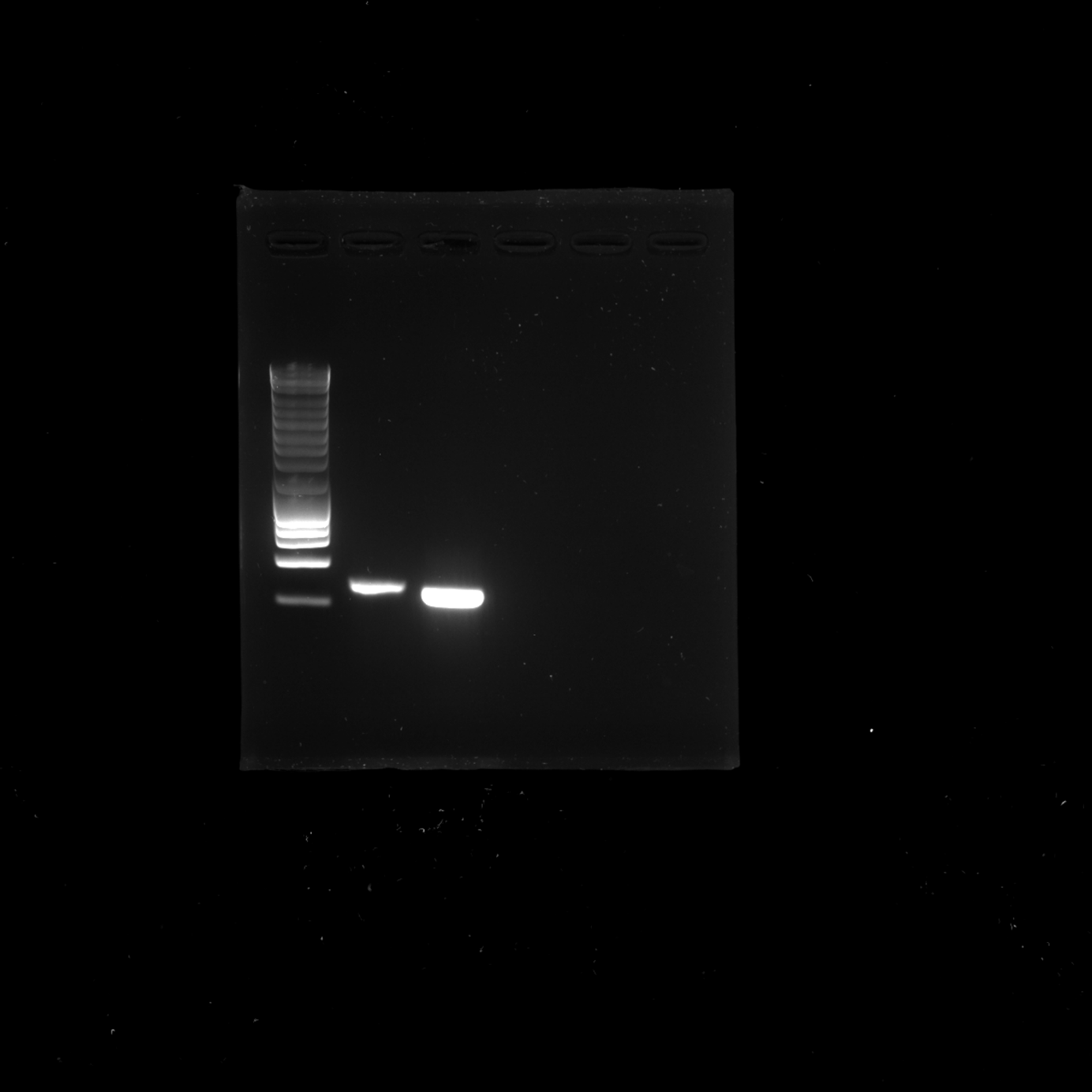

Figure S6. The result of *PaCYP7A1* *in vitro* transcription reaction. The lane 1 is the Eurogentec 1 Kb SamrtLadder. Lane 2 is the PCR product of the *PaCYP7A1* used as a control. Lane 3 is the purified dsCYP7 produced via *in vitro* transcription.

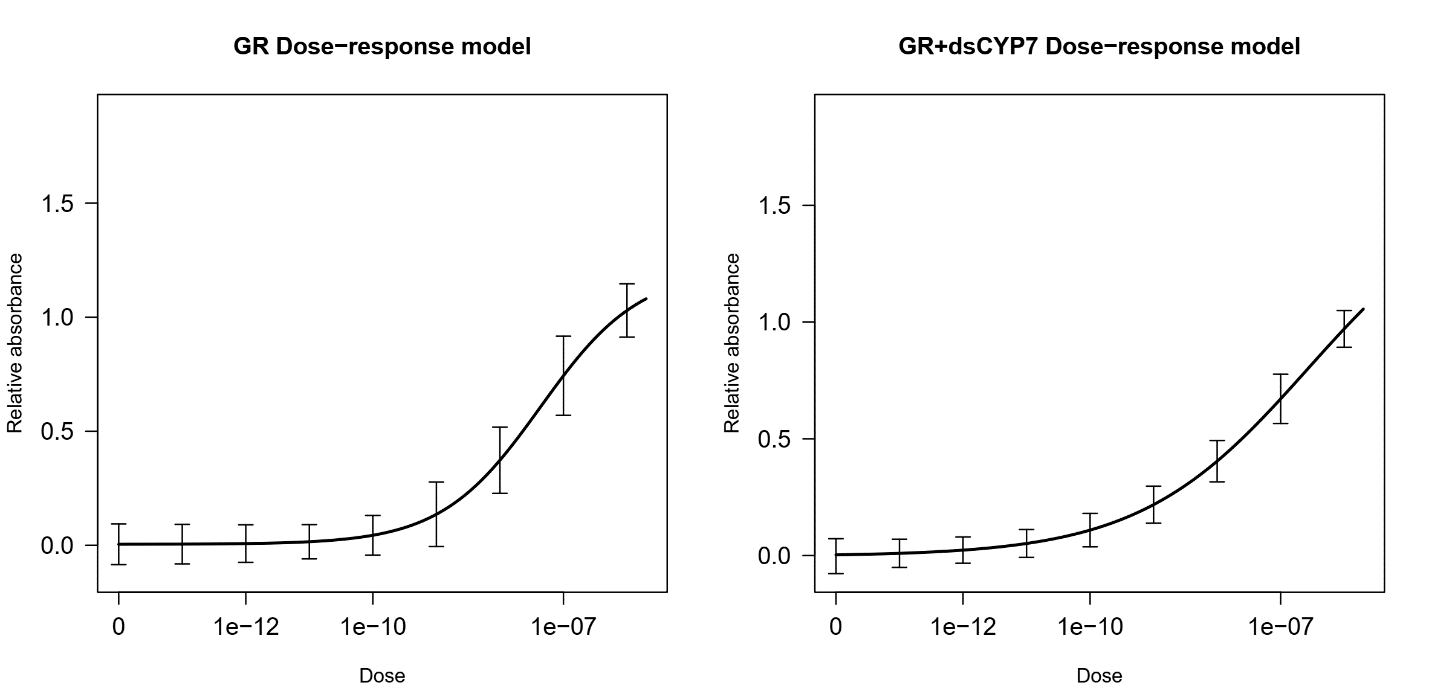

Figure S7. The dose-response curve obtained from absorbance reading data in the experiment with 0.7 µM dsCYP7.

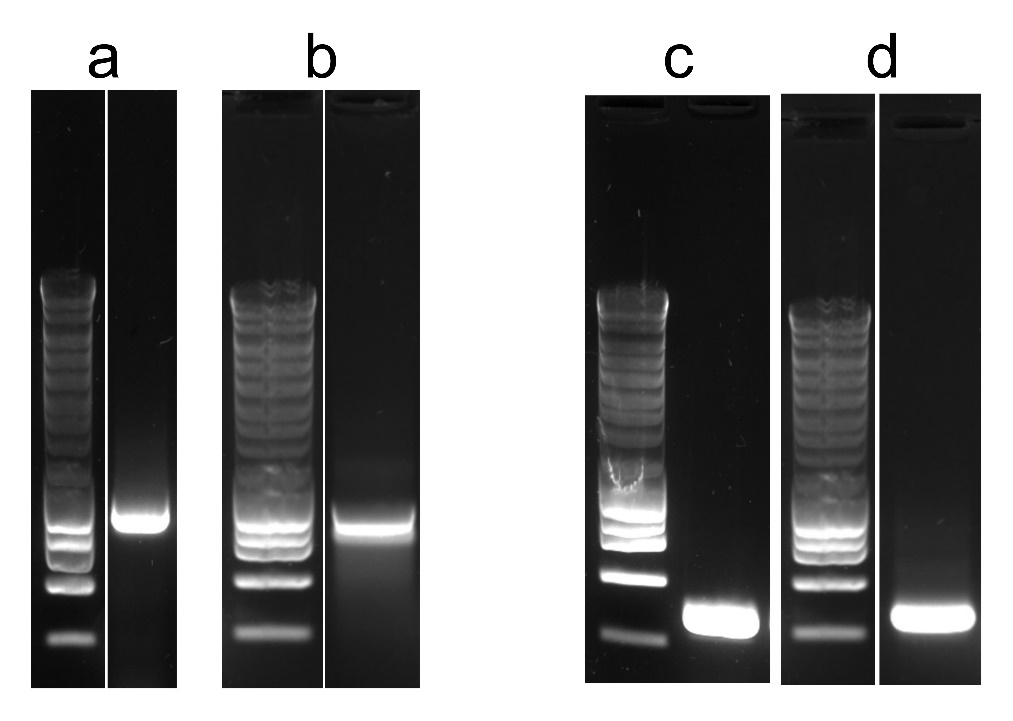

Figure S8. The result of *in vitro* transcription reactions for dsM6PR and dsGFP. A) PCR product of M6PR fragment. B) dsM6PR. C) PCR product of GFP fragment. D) dsGFP. In all photos, lane 1 is the Eurogentec 1 Kb SamrtLadder.

A

B

D

C

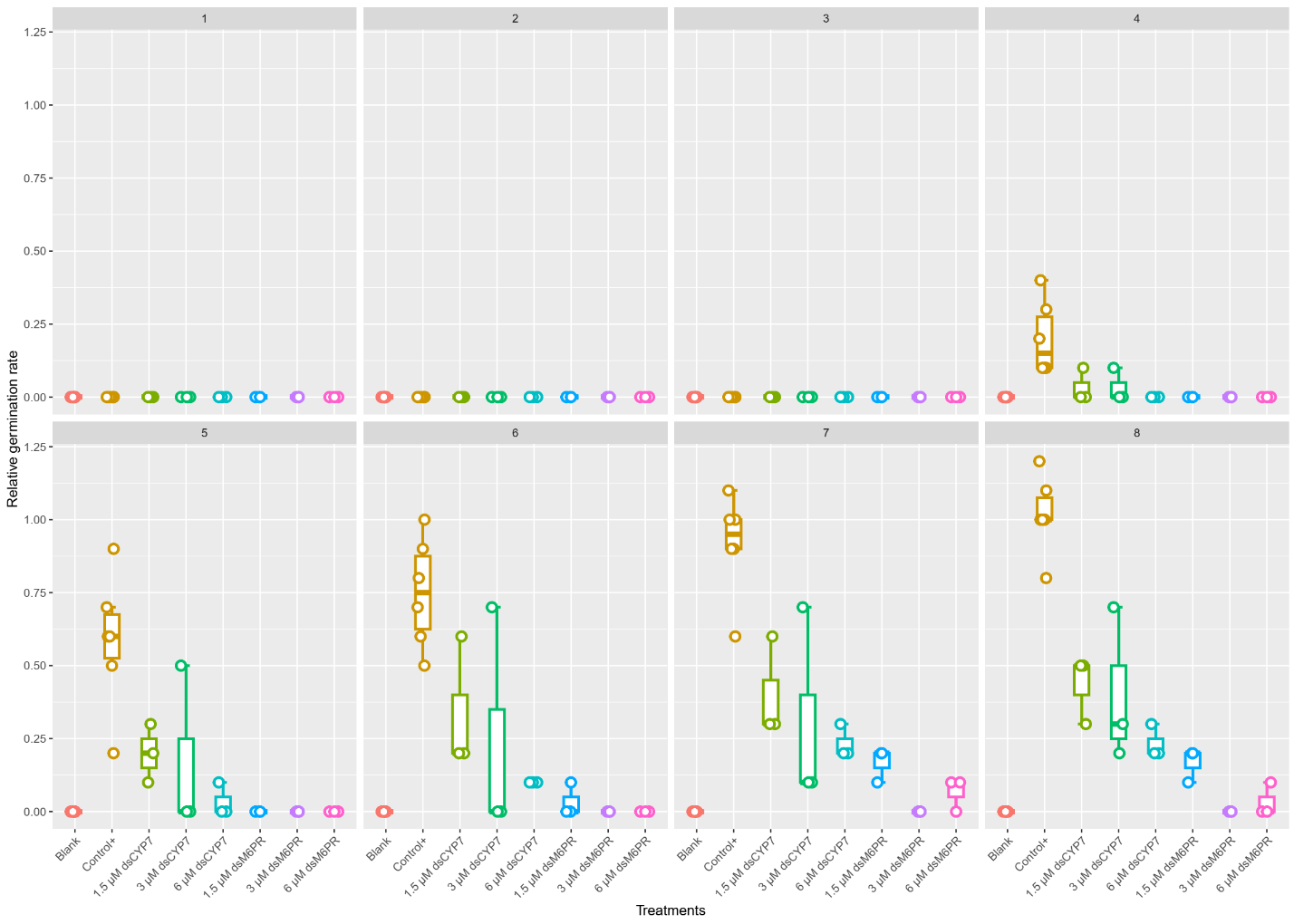

Figure S9. The boxplot of the relative germination rates obtained in the experiment with dsCYP7 as well as dsM6PR as a non-specific dsRNA from 1- to 8-dpt. Seeds were treated with 6, 3, and 1.5 µM of each dsRNA. 10^-6^ M (+)GR24 was considered as the positive control and seeds without (+)GR24 were considered as blank.

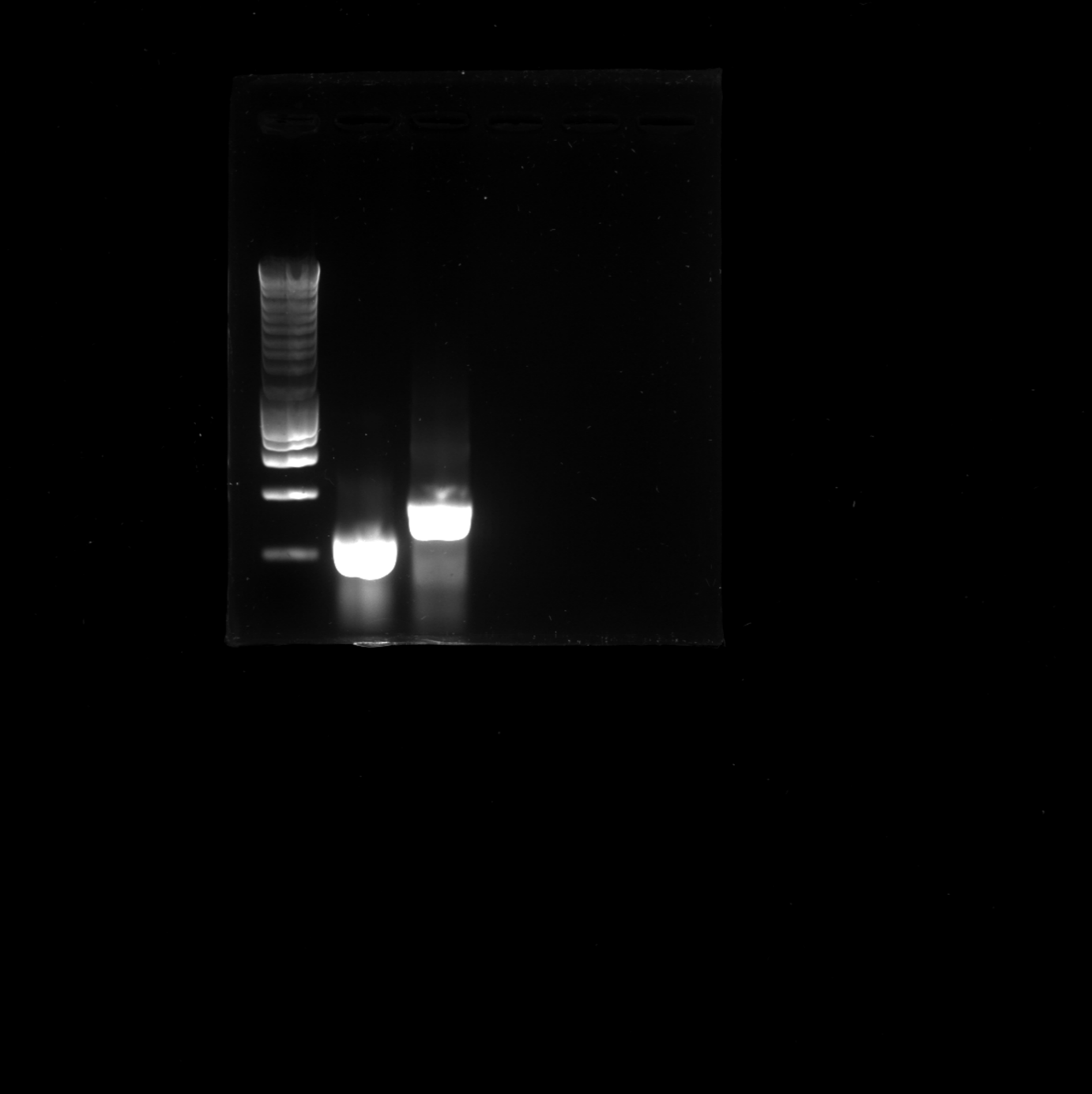

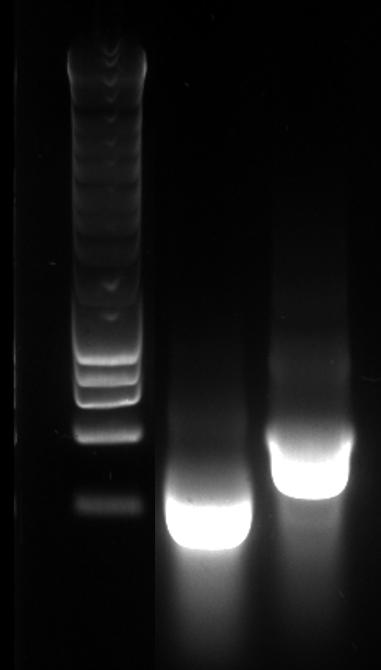

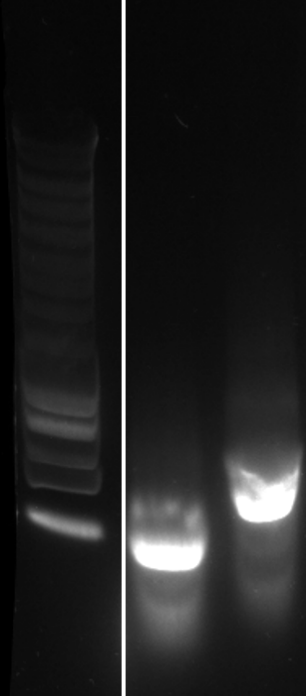

Figure S10. dsCYP7 and dsM6PR obtained from the seed plate. Lane 1-3 are 1 kb Eurogentec DNA SmartLadder, dsCYP7, and dsM6PR, respectively. A) before precipitation, B) after precipitation with ammonium acetate, and C) after purification with MEGAclear.

A

B

C
